# Intrinsic Excitability Increase in Cerebellar Purkinje Cells Following Delay Eyeblink Conditioning in Mice

**DOI:** 10.1101/306639

**Authors:** Heather K. Titley, Gabrielle V. Watkins, Carmen Lin, Craig Weiss, Michael McCarthy, John F. Disterhoft, Christian Hansel

## Abstract

Cerebellar learning is canonically thought to rely on synaptic plasticity, particularly at synaptic inputs to Purkinje cells. Recently, however, other complementary mechanisms have been identified. Intrinsic plasticity is one such mechanism, and depends in part on the down-regulation of calcium-dependent SK-type K channels, which is associated with an increase in neuronal excitability. In the hippocampus, SK-mediated intrinsic plasticity has been shown to play a role in trace eyeblink conditioning; however, it is not yet known how intrinsic plasticity contributes to a cerebellar learning task such as delay eyeblink conditioning. Whole cell recordings were obtained from acute cerebellar slices from mice ~48 hours after learning a delay eyeblink conditioning task. Over a period of repeated training sessions mice received either distinctly paired trials of a tone co-terminating with a periorbital shock (conditioned mice) or trials in which these stimuli were presented in an unpaired manner (pseudoconditioned mice). Conditioned mice show a significantly reduced afterhyperpolarization (AHP) following trains of parallel fiber stimuli. Moreover, we find that SK-dependent intrinsic plasticity is occluded in conditioned, but not pseudoconditioned mice. These findings show that excitability is enhanced in Purkinje cells after delay eyeblink conditioning and point toward a downregulation of SK channels as a potential underlying mechanism.

## Introduction

Classically, cerebellar motor learning has been thought to rely on changes in synaptic strength, specifically at synapses between parallel fibers (PFs) and Purkinje cells in the cerebellar cortex (for review, see Jörntell and Hansel, 2006). It has been suggested that cerebellar learning may also involve other non-synaptic (‘intrinsic’) plasticity mechanisms (see Titley et al., 2017). While excitability changes have indeed been demonstrated subsequent to delay eyeblink conditioning (Schreurs et al., 1998), this surprising result (excitability changes were observed in an average measure across ALL recorded Purkinje cells from conditioned rabbits) has not yet been independently confirmed, and the phenomenon has not been studied in further detail.

In cerebellar Purkinje cells, a form of intrinsic plasticity that is mediated by SK2 channel down-regulation (Purkinje cells only express the SK2 isoform; Cingolani et al., 2002) results in enhanced excitability, enhanced spine calcium signaling, and a lower probability for subsequent LTP induction (Belmeguenai et al., 2010; Hosy et al., 2011; note that in Purkinje cells the calcium threshold for LTD is higher than that for LTP: Coesmans et al., 2004; Piochon et al., 2016). SK channels contribute to an afterhyperpolarization (AHP) following bursts of action potentials and are involved in the regulation of spike firing frequency in some neurons (Stocker et al., 1999; Pedarzani et al., 2001). In the cerebellum, SK2 channels are known to be involved in the medium-slow AHP mediating the complex spike pause in Purkinje cells (Kakizawa et al., 2007). Indeed, intrinsic plasticity reduces the duration of complex spike pauses in Purkinje cells and this type of plasticity depends on SK channel modulation (Grasselli et al., 2016).

In the hippocampus, SK2 is known to be important for synaptic plasticity and learning. SK2 overexpression reduced long-term potentiation in hippocampal slices, and severely impaired hippocampal dependent learning in mice (Hammond et al., 2006). During trace eyeblink conditioning in rats, an SK channel activator reduced the excitability of CA1 pyramidal neurons by increasing the AHP, and impaired learning (McKay et al., 2012, 2013). Here, we ask whether intrinsic excitability is changed in a cerebellar motor learning task (delay eyeblink conditioning), and whether the altered excitability parameters point toward an involvement of SK channel modulation.

## Methods

### Animals

All procedures were approved by and performed in accordance with the guidelines of the Animal Care and Use Committees of both Northwestern University and the University of Chicago. Experiments were performed using 5-8 week old male mice (C57BL/6J) obtained from Jackson Laboratory.

### Surgery

Briefly, animals were anesthetized with 3-4% vaporized isoflurane mixed with oxygen at a flow rate of 1-2 liters per minute, with buprenorphine (0.05—2 mg/kg) given subcutaneously as an analgesic. Mice were placed in a stereotaxic device and a midline incision was made along the scalp. The skin was retracted laterally and the periosteum was scraped away. Two small screws (one in front of Bregma and one in front of Lambda) were implanted into the skull left of the midline. A ground wire of a headpiece containing five wires (one ground, two shock-delivering, and two EMG recording wires) was wrapped around the screws in a figure eight pattern. A thin layer of adhesive cement was placed on the skull, the screws, and the wire. The skin around the right eye was retracted to reveal the muscle around the eye. Two shock-delivering wires were placed under the skin, caudal to the right eye and two EMG recording wires were placed on the musculus orbicularis behind the eyelid. The base of the wires was stabilized with another layer of cement. The headpiece was stabilized with dental cement. The animal was allowed to recover on a warmed heating pad before the animal was placed back in its home cage. Mice were allowed to recover one week after the surgery before starting eyeblink conditioning.

### Delay Eyeblink Conditioning

Animals were randomly assigned to either a conditioned group or a pseudoconditioned group. The training apparatus included a cylindrical treadmill inside a sound attenuating training chamber (IAC Acoustics). Mice were head fixed and allowed to run freely on the cylindrical treadmill (see Lin et al., 2016). Mice were allowed to habituate to the chamber and cylindrical treadmill during two separate sessions the day before training. During each habituation session, the mice were placed on the cylindrical treadmill for the same duration as one training session, although no stimuli were presented. Training began the day following habituation. The mice were trained for two sessions a day for three days, with a two hour interval between training sessions. Conditioned mice were given the delay eyeblink conditioning paradigm, which consisted of a 75 dB tone (350 ms, 2 kHz) conditioned stimulus (CS) paired with a 100 ms unconditioned stimulus (US) consisting of 6 sets of biphasic shocks (120 Hz; 1 ms/pulse) to the periorbital region using biphasic stimulus isolator (WPI model A385). The US intensity was adjusted for each mouse to elicit a reliable blink (0.3-2 mA). The US overlapped with and coterminated with the CS. Each conditioning session consisted of 30 paired CS-US trials with a random 30-60 second inter-trial interval. The pseudoconditioned mice were presented with 30 unpaired CS trials and 30 unpaired US trials in pseudorandomized order with a 15-30 second inter-trial interval.

EMG signals were amplified and filtered through 100 to 5 kHz low/high pass filter. The signals were rectified and integrated with a time constant of 10 ms. Baseline EMG activity was defined as the average activity 250 ms before CS onset. Conditioned responses were identified as having EMG activity 4 standard deviations above the baseline, present at least 20 ms before US onset. The electrophysiological experiments described below were performed with the experimenter blind to the training condition of each mouse.

### Slice Preparation

Within ~48 hours of the final delay eyeblink conditioning training session, animals were anaesthetized with isoflurane and immediately decapitated. The cerebellum was removed and the right paravermis / hemisphere isolated in artificial cerebrospinal fluid (ACSF) cooled to 1-4º C, and containing (in mM): 124 NaCl, 5 KCl, 1.25 Na_2_HPO_4_, 2 CaCl_2_, 2 MgSO_4_, 26 NaHCO_3_, and 10 D-glucose, bubbled with 95% O_2_ and 5% CO_2_. Parasagittal slices of the right hemisphere and paravermis (200μm) containing lobule HVI were prepared with a Leica VT-1000S vibratome, then incubated for at least 1 hr at room temperature in oxygenated ACSF.

### Somatic Whole-Cell Patch-Clamp Recordings

Slices were continuously perfused with ACSF containing picrotoxin (100 μM, Sigma Aldrich) to block GABA_A_ receptors, and held at near physiological temperature (32-34°C) over the course of the experiments. Slices were visualized using an x40 objective mounted on either a Zeiss Examiner A1 microscope (Carl Zeiss MicroImaging) or a Zeiss Axioskop 2 FS plus microscope (Carl Zeiss MicroImaging). Patch-clamp recordings were made from Purkinje cells located at the base of the primary fissure using an EPC-10 amplifier (HEKA Electronics). Currents were filtered at 3kHz, sampled at 20kHz, and acquired using Patchmaster software (HEKA Electronics). The access resistance was compensated (70-80 %) in current clamp mode. In most experiments hyperpolarizing bias currents were applied to hold the membrane potential around −70 mV. Patch pipettes (2-6 MΩ) were filled with internal solution containing (in mM): 120 K-gluconate, 9 KCl, 10 KOH, 3.48 MgCl_2_, 10 HEPES, 4 NaCl, 4 Na_2_ATP, 0.4 Na_3_GTP, and 17.5 sucrose, with the pH adjusted to 7.25-7.35.

Spontaneous activity was determined by measuring cell firing in current-clamp mode over 1-second sweeps; Purkinje cells that did not generate action potentials with 0 pA injected current (no bias current) were excluded from the experiments. Evoked activity was measured by holding Purkinje cells at −70 mV in current-clamp mode and applying a 500 msec depolarization step that gradually increased in 50 pA steps. Parallel fibers were stimulated by placing a glass electrode filled with ACSF in the molecular layer around the distal dendrites of the Purkinje cell while held at −70 mV. Parallel fiber inputs were stimulated 10 times at 100 Hz.

Intrinsic plasticity was monitored during the test periods by injecting a 500 msec depolarizing current (300-800 pA) to evoke 10-15 action potentials. The input resistance was monitored throughout the experiments by applying hyperpolarizing current steps (−100 pA) at the end of each sweep. During tetanization, depolarizing currents (100 ms) were injected at 5Hz for 8 sec. Intrinsic plasticity recordings were excluded if the input resistance (R_i_) or holding potential (V_h_) changed by ≥15% over the course of the baseline.

### Data Analysis

CS and US stimuli were delivered and EMG signals were collected using custom software written for LabVIEW (National Instruments). Cellular data were analyzed using Excel (Microsoft), Igor (Wave-metrics), and Statistica (Tibco). All data are expressed as mean ± SEM. Statistical analyses were performed using the Mann-Whitney U test (between groups), paired Student's t-test (within subjects), and two-way repeated measures ANOVA tests as appropriate.

## Results

### Mice successfully learned the associative conditioning task

As expected, mice successfully learned to associate an auditory stimulus (conditioned stimulus, CS) with a periorbital shock to the eyelid (unconditioned stimulus, US) when the CS preceded and co-terminated with the US. Over six consecutive sessions, conditioned mice learned to blink their eye in response to the tone before the actual US was presented (conditioned response, CR). Figure 1A shows the gradual acquisition of the CR in the conditioned mice (green dots, n=37) and the absence of a similar learned response in pseudoconditioned mice (black dots, n=27). While many mice were trained, we later imposed a learning criterion to ensure that cell physiological parameters were only collected from successfully trained conditioned mice or pseudoconditoned mice. As a result we eliminated conditioned mice that failed to reach at least 80% CR during any session (11 mice), as well as any pseudoconditioned mice that reached ≥ 50% CR (6 mice). Furthermore, due to our strict location cell specificity, recordings were not obtained from all mice, and thus we eliminated mice from the curve from which we did not obtain cellular data (13 conditioned mice, 8 pseudoconditioned mice).

**Figure 1.**
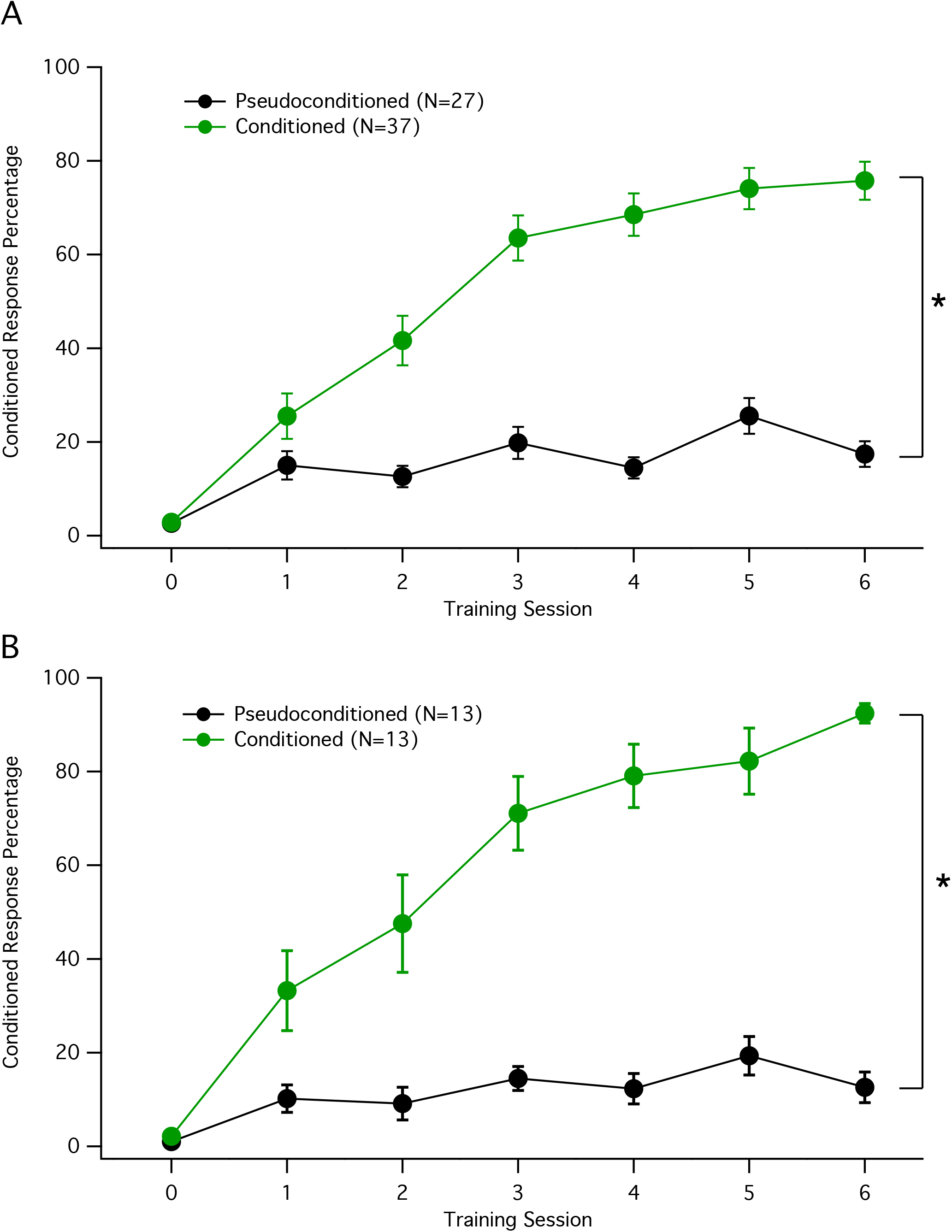
Delay eyeblink conditioning generates learned behavior. Mice were divided into two groups: conditioned animals that received paired CS (tone) and US (shock) stimuli (green), and pseudoconditioned animals that received unpaired stimuli over the same number of trials (black). Conditioned animals were successfully able to learn and generate a conditioned response (CR, eye closure) at a high percentage by the end of the sixth learning session. **A.** Data from all mice trained. **B.** Learning curves from the subset of mice that met the learning criterion and from which cells were successfully obtained. Data are presented as mean ± SEM. * Indicates significance of p<0.05.

Figure 1B shows the remaining learning curves from mice that met the learning criterion and from which we obtained cellular data. After the 6^th^ learning session conditioned mice (n=13) were able to show a CR 92.4 ± 2.1 % of the trials. In contrast, the pseudoconditioned mice (n=13) that received unpaired stimuli of the tone and the shock showed a CR response rate of 12.6 ± 3.3 %. Therefore conditioned animals acquired a significantly greater percent CR over the course of the training trials relative to the pseudoconditioned animals (two-factor repeated measures ANOVA, F(6,168)=13.03, p<0.001).

### Spontaneous and evoked spiking activity were not affected by associative learning

To evaluate potential cellular differences in cerebellar Purkinje cells caused by conditioning, conditioned and pseudoconditioned mice were sacrificed ~48 hours after the last training trial. We examined Purkinje cells by whole-cell patch clamp electrophysiology deep in the primary fissure near cerebellar lobules V/HVI, which has been shown to be the microzone involved with delay eyeblink conditioning (Heiney et al., 2014; see also Mostofi et al., 2010; Steinmetz and Freeman, 2014). The experimenters were blind to the condition of the mouse during the time of the experiment and analysis. To ensure that the Purkinje cells examined were healthy and displayed normal physiological properties, only cells that produced spontaneous action potentials without bias current injection were analyzed.

To determine whether delay eyeblink conditioning increased spike firing of Purkinje cells, we measured both spontaneous and evoked firing under current clamp conditions. The eyeblink conditioning task had no effect on the spontaneous firing frequency of cells. Figure 2B illustrates example traces from two Purkinje cells firing spontaneously. The spontaneous firing frequency recorded from Purkinje cells in pseudoconditioned mice was 73.4 ± 6.0 spikes/s (n=20). This was not significantly different from Purkinje cells from conditioned mice with a spontaneous firing rate of 81.5 ± 7.8 Hz (n=17; Mann-Whitney U test, p=0.43; Figure 2A).

**Figure 2.**
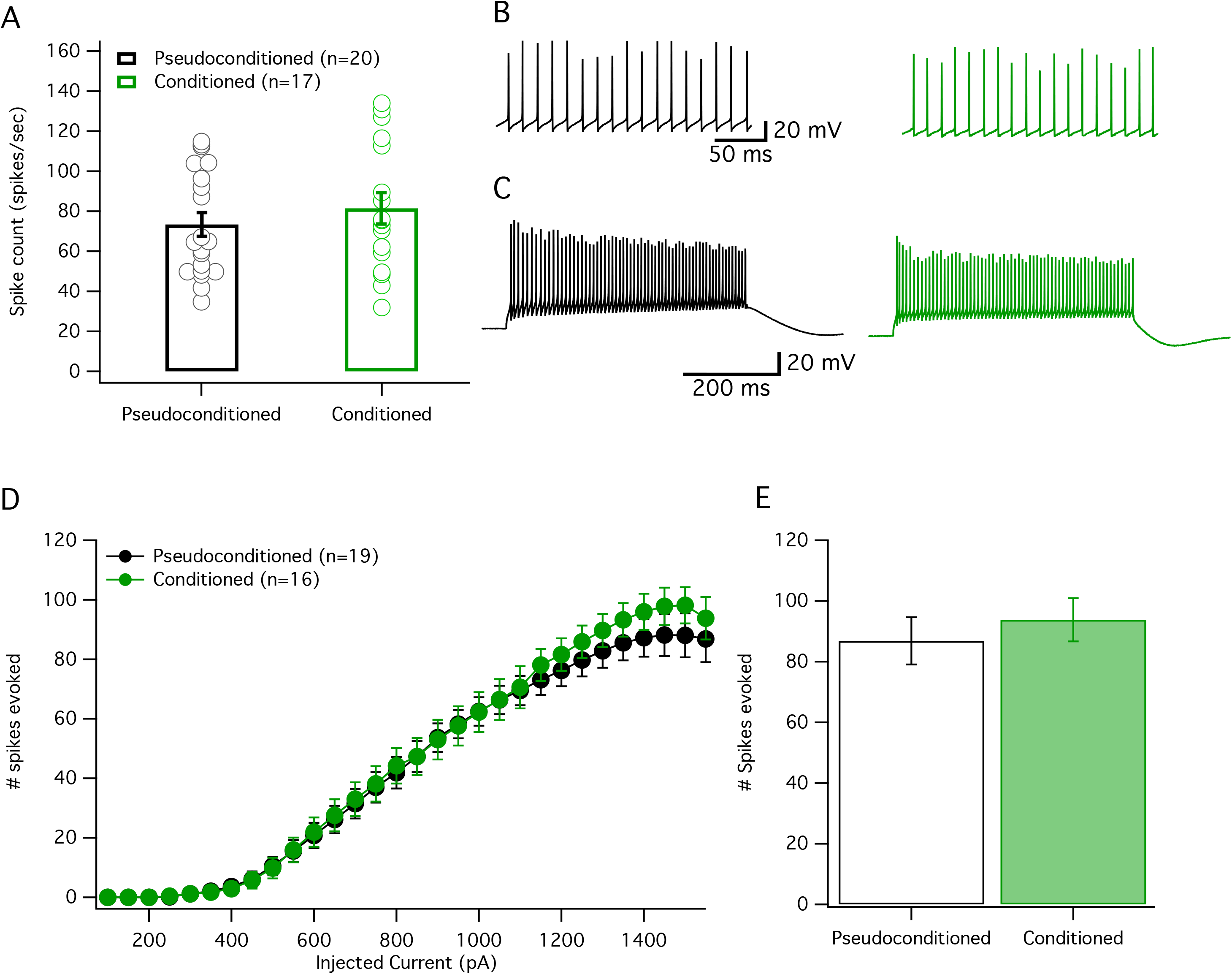
Delay eyeblink conditioning has no effect on spontaneous or evoked activity of cerebellar Purkinje cells. **A.** Spike count of action potentials per second measured from spontaneously firing cells (with no injected currents) from conditioned (green) and pseudoconditioned mice (black). Dots represent individual cells, while bars represent the mean average. **B.** Example traces depicting spontaneous firing from cells in panel A. **C.** Example traces depicting evoked firing from cells in panels D and E with 1550 pA current injected. **D.** Mean evoked firing from conditioned (green) and pseudoconditioned cells (black) over a broad range of injected currents. **E.** The mean evoked number of spikes produced when the injected current is 1550 pA in the conditioned and pseudoconditioned cells taken from panel D. Data are presented as mean ± SEM.

To further analyze the excitability of Purkinje cells we looked at firing evoked by injection of increasing depolarization steps. Under current clamp conditions Purkinje cells were held at a potential of around −70mV, while depolarizing currents (500 ms, 100-1550 pA) were injected into the cell in 50 pA steps to induce firing. We analyzed the curves of spikes evoked plotted against injected current. Figure 2C shows that the curves by the conditioned and pseudoconditioned cells appeared very similar and were not significantly different (two-way repeated measures ANOVA, F(26, 832)=0.81, p=0.74). When 1550 pA of current was injected into the cells, no differences were found between the number of spikes evoked from cells of conditioned (93.8 ± 7.1 spikes; n=16) or pseudoconditioned mice (86.8 ± 7.8 spikes; n=19; Mann-Whitney U test, p=0.61; Figure 2D). Together, these results suggest that delay eyeblink conditioning did not affect spontaneous or depolarization-evoked spike firing.

### Decreased amplitudes of AHPs following parallel fiber bursts in conditioned mice

As SK2 channels have been shown to underlie the medium AHP in Purkinje cells (Kakizawa et al., 2007; Grasselli et al., 2016), we looked at the AHP following a burst of 10 parallel fiber stimuli at 100 Hz. Figure 3 shows example traces for Purkinje cells from a pseudoconditioned (Figure 3A) and a conditioned animal (Figure 3B) following a burst of parallel fiber stimulation. Cells from the conditioned mice were found to have a significantly reduced AHP compared to cells from pseudoconditioned animals. The average minimum (peak) amplitude of the AHP was significantly lower in cells from conditioned mice (−4.6 ± 0.5 mV; n=12) than in cells from pseudoconditioned mice (−6.1 ± 0.5 mV; n=12; Mann-Whitney U test, p=0.04; Fig. 3C). Furthermore, in the same recordings we found a significant reduction in the total AHP area (conditioned: −455.3 ± 46.7 ms*mV; pseudoconditioned: −709.8 ± 50.1 ms*mV; Mann-Whitney U test, p=0.012; Figure 3D) as well as a reduction in AHP duration (conditioned: 206.5 ± 13.2 ms; pseudoconditioned: 269.2 ± 18.0 ms; Mann-Whitney U test, p=0.001; Figure 3E). Therefore, Purkinje cells in mice that received delay eyeblink conditioning showed a significantly reduced AHP compared to Purkinje cells from pseudoconditioned control mice, which is compatible with a downregulation of SK2 channels during eyeblink conditioning.

**Figure 3.**
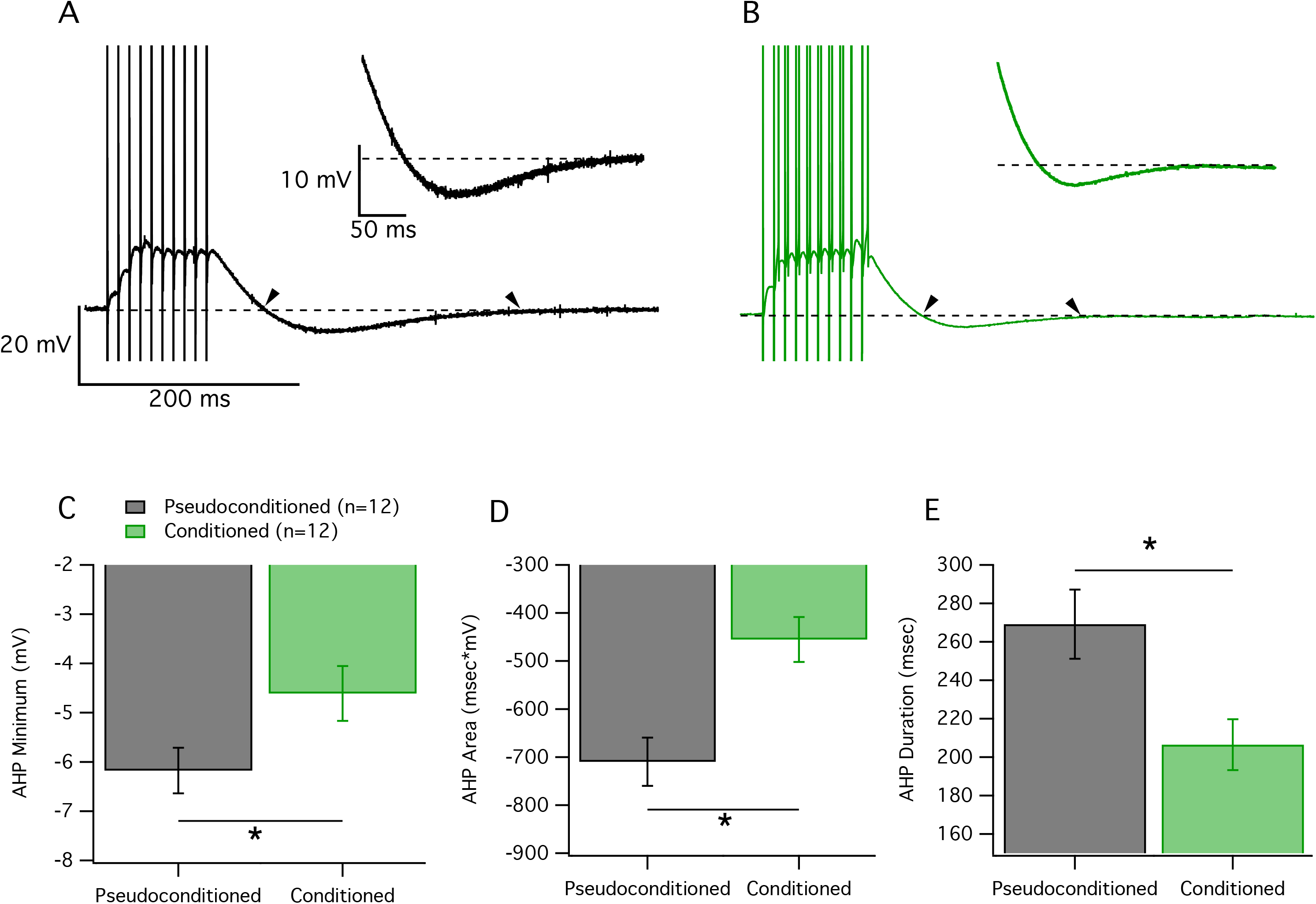
Eyeblink conditioning reduced the AHP following a burst of parallel fiber stimulation. **A and B.** Example traces of the response to 10 parallel fiber stimuli at 100 Hz from cells from a pseudoconditioned mouse (A) and a conditioned mouse (B). Dashed line indicates the baseline potential at −70 mV. Arrowheads indicate the start and finish of the AHP. Inset: A close up showing the AHP at higher magnification. **C-E.** Measures of the AHP following the parallel fiber burst in cells from conditioned (green) and pseudoconditioned mice (black). **C.** Minimum amplitude of AHP. **D.** Area of AHP (measured as area between curve and baseline). **E.** Duration of AHP. Data are presented as mean ± SEM. * Indicates significance of p<0.05.

### Intrinsic plasticity is occluded in cells from conditioned mice

To further test the involvement of SK2 in eyeblink conditioning, we repeatedly depolarized (5 Hz, 8 sec) the Purkinje cells to induce intrinsic plasticity (Belmeguenai et al., 2010). Pseudoconditioned mice showed a significant increase in the number of spikes evoked following the intrinsic potentiation protocol compared to baseline (paired Student's t-test, p=0.004; Figure 4). In contrast, Purkinje cells from conditioned mice failed to show increases in excitability that were significantly different from baseline (paired Student's t-test, p=0.08). When comparing the difference between these groups at 21-25 mins after tetanization, Purkinje cells from the pseudoconditioned group showed significantly larger excitability changes compared to cells from conditioned mice (pseudoconditioned: 146.0 ± 9.9 %; n=9; conditioned: 112.4 ± 6.4 %; n=11; Mann-Whitney U test, p=0.007). Therefore, conditioned mice failed to show significant intrinsic potentiation following tetanization, while pseudoconditioned mice were able to upregulate excitability, suggesting that SK2-dependent intrinsic plasticity was occluded after delay eyeblink conditioning.

**Figure 4.**
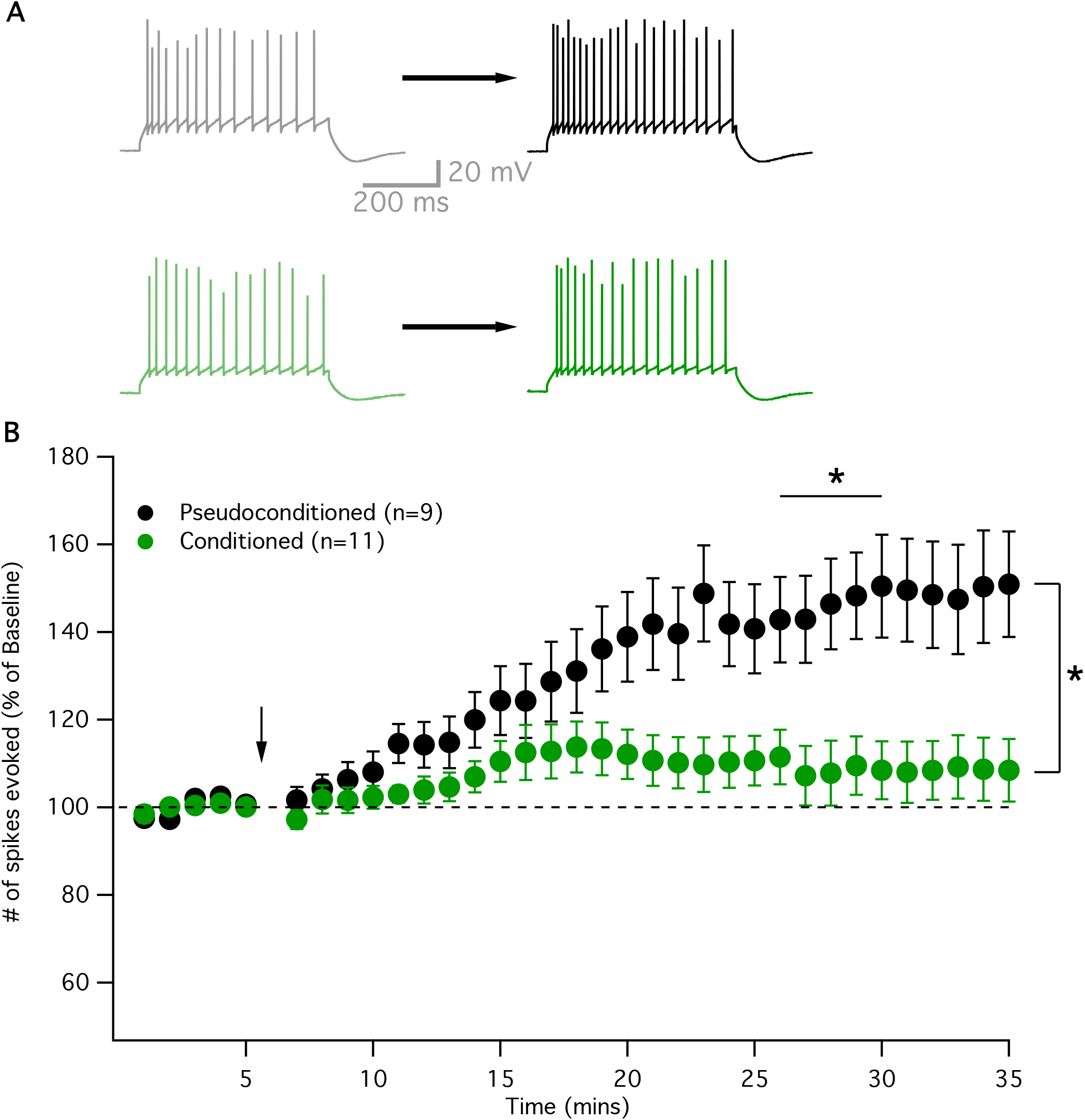
Intrinsic plasticity is occluded in Purkinje cells from conditioned animals. **A.** Example traces of evoked activity during the baseline (left) and following tetanization (right) in pseudoconditoned (black) and conditioned (green) mice. **B.** Time graph showing intrinsic potentiation (increase in evoked spiking as percentage of baseline) in pseudoconditioned cells (black) following tetanization (arrow), but not in conditioned cells (green). Values are presented as mean ± SEM. * Indicates significance of p<0.05.

## Discussion

### AHP is reduced in Purkinje cells after delay eyeblink conditioning

It has previously been reported that the AHP amplitude is reduced in rats after trace eyeblink conditioning tasks in both CA1 and CA3 pyramidal neurons (Moyer et al., 1996; Thompson et al., 1996). Similarly, Schreurs and colleagues found a reduction in the AHP in Purkinje cells from conditioned rabbits compared to cells from pseudoconditioned animals in a delay eyeblink conditioning task. In their study they found that Purkinje cells in area HVI from conditioned animals showed a decrease in dendritic spike threshold as well as a reduction in the AHP following depolarization steps which persisted even one month after the final training session (Schreurs et al., 1998). Here, we similarly found that after a burst of parallel fiber stimulation, Purkinje cells in mice that were conditioned showed a significantly reduced AHP minimum amplitude, AHP duration, as well as a reduced AHP area (Figure 3). The AHP is known to be involved in spike excitability, and in Purkinje cells it is known to be partially mediated by SK2 conductances, suggesting that SK channels might be down-regulated during a delay eyeblink conditioning task.

### Intrinsic potentiation is occluded after delay eyeblink conditioning

To further test for the involvement of SK channels, we tetanized cells using an intrinsic plasticity protocol. In Purkinje cells, this form of intrinsic plasticity is known to depend on SK2 channels, as SK2 is the only isoform expressed in Purkinje cells. Intrinsic potentiation is blocked by the SK inhibitor apamin (Ohtsuki et al., 2012) and is shown to be absent in SK2 knockout mice (Grasselli et al., 2016). Here, we show that conditioned mice failed to increase their spike firing following an intrinsic potentiation protocol, while pseudoconditioned cells were able to upregulate evoked spike firing (Figure 4). This suggests that intrinsic plasticity was occluded in the conditioned mice, suggesting that this non-synaptic form of plasticity is involved in learning (Titley et al., 2017). Here, we report the effects in slices 48 hours after learning has ended, although the maximal duration of this effect is not known.

Together, it seems delay eyeblink conditioning is associated in Purkinje cells with intrinsic plasticity as reflected by ta reduction in the AHP. We further suggest that the resulting increase in membrane excitability is a consequence of a downregulation of SK2 conductances, which provide a critical component of the medium AHP, and are involved in the form of intrinsic plasticity tested here.

## Acknowledgments

• This study was supported by a grant from the National Institute of Neurological Disorders and Stroke (NS-062271 to C.H.) and from the National Institute on Aging (R37 AG008796 to J.F.D and F31AG055331 to C.L.). The authors would like to thank members of the Hansel and Disterhoft labs for many helpful discussions.

## References

Belmeguenai A, Hosy E, Bengtsson F, Pedroarena CM, Piochon C, Teuling E, He Q, Ohtsuki G, De Jeu MTG, Elgersma Y, De Zeeuw CI, Jörntell H, Hansel C (2010) Intrinsic plasticity complements long-term potentiation in parallel fiber input gain control in cerebellar Purkinje cells. J Neurosci 30: 13630–13643.

Coesmans M, Weber JT, De Zeeuw CI, Hansel C (2004) Bidirectional parallel fiber plasticity in the cerebellum under climbing fiber control. Neuron 44:691–700.

Grasselli G, He Q, Wan V, Adelman JP, Ohtsuki G, Hansel C (2016) Activity-Dependent Plasticity of Spike Pauses in Cerebellar Purkinje Cells. Cell Rep 14:2546–2553.

Hammond RS, Bond CT, Strassmaier T, Ngo-Anh TJ, Adelman JP, Maylie J, Stackman RW (2006) Small-conductance Ca2+-activated K+ channel type 2 (SK2) modulates hippocampal learning, memory, and synaptic plasticity. J Neurosci 26:1844–1853.

Heiney SA, Kim J, Augustine GJ, Medina JF (2014) Precise control of movement kinematics by optogenetic inhibition of Purkinje cell activity. J Neurosci 34: 2321–2330.

Hosy E, Piochon C, Teuling E, Rinaldo L, Hansel C (2011) SK2 channel expression and function in cerebellar Purkinje cells. J Physiol 589:3433–3440.

Jörntell H, Hansel C (2006) Synaptic memories upside down: bidirectional plasticity at cerebellar parallel fiber-Purkinje cell synapses. Neuron 52:227–238.

Kakizawa S, Kishimoto Y, Hashimoto K, Miyazaki T, Furutani K, Shimizu H, Fukaya M, Nishi M, Sakagami H, Ikeda A, Kondo H, Kano M, Watanabe M, Iino M,Takeshima H (2007) Junctophilin-mediated channel crosstalk essential for cerebellar synaptic plasticity. EMBO J 26:1924–1933.

Lin C, Disterhoft J, Weiss C (2016) Whisker-signaled Eyeblink Classical Conditioning in Head-fixed Mice. J Vis Exp 109.

McKay BM, Oh MM, Disterhoft JF (2013) Learning Increases Intrinsic Excitability of Hippocampal Interneurons. J Neurosci 33:5499–5506.

McKay BM, Oh MM, Galvez R, Burgdorf J, Kroes RA, Weiss C, Adelman JP, Moskal JR, Disterhoft JF (2012) Increasing SK2 channel activity impairs associative learning. J Neurophysiol 108:863–870.

Mostofi A, Holtzman T, Grout AS, Yeo CH, Edgley SA (2010) Electrophysiological localization of eyeblink-related microzones in rabbit cerebellar cortex. J Neurosci 30:8920–8934.

Moyer JR, Thompson LT, Disterhoft JF (1996) Trace eyeblink conditioning increases CA1 excitability in a transient and learning-specific manner. J Neurosci 16:5536–5546.

Ohtsuki G, Piochon C, Adelman JP, Hansel C (2012) SK2 channel modulation contributes to compartment-specific dendritic plasticity in cerebellar Purkinje cells. Neuron 75:108–120.

Pedarzani P, Mosbacher J, Rivard A, Cingolani LA, Oliver D, Stocker M, Adelman JP, Fakler B (2001) Control of electrical activity in central neurons by modulating the gating of small conductance Ca2+-activated K+ channels. J Biol Chem.

Piochon C, Titley HK, Simmons DH, Grasselli G, Elgersma Y, Hansel C (2016) Calcium threshold shift enables frequency-independent control of plasticity by an instructive signal. Proc Natl Acad Sci 113:13221–13226.

Schreurs BG, Gusev PA, Tomsic D, Alkon DL, Shi T (1998) Intracellular correlates of acquisition and long-term memory of classical conditioning in Purkinje cell dendrites in slices of rabbit cerebellar lobule HVI. J Neurosci 18:5498–5507.

Steinmetz AB, Freeman JH (2014) Localization of the cerebellar cortical zone mediating acquisition of eyeblink conditioning in rats. Neurobiol Learn Mem 114:148–154.

Stocker M, Krause M, Pedarzani P (1999) An apamin-sensitive Ca2+-activated K+ current in hippocampal pyramidal neurons. Proc Natl Acad Sci U S A.

Thompson LT, Moyer JR, Disterhoft JF (1996) Transient changes in excitability of rabbit CA3 neurons with a time course appropriate to support memory consolidation. J Neurophysiol 76:1836–1849.

Titley HK, Brunel N, Hansel C (2017) Toward a Neurocentric View of Learning. Neuron 95:19–32.

